# Ethanol induces craniofacial defects in Bmp mutants independent of *nkx2.3* by elevating cranial neural crest cell apoptosis

**DOI:** 10.1101/2024.12.31.630963

**Authors:** Hieu Vo, C. Ben Lovely

## Abstract

**Background:** Fetal Alcohol Spectrum Disorders (FASD) describes a wide range of neurological defects and craniofacial malformations associated with prenatal ethanol exposure. While there is growing evidence for a genetic component to FASD, little is known of the cellular mechanisms underlying these ethanol-sensitive loci in facial development. Endoderm morphogenesis to form lateral protrusions called pouches is one key mechanism in facial development. We have previously shown that multiple members of the Bone Morphogenetic Pathway (Bmp) signaling pathway, a key regulator of pouch formation, interacts with ethanol disrupting facial development. However, ethanol does not directly impact Bmp signaling suggesting that downstream effectors, like *nkx2.3* may mediate the impact of ethanol on Bmp mutants.

**Methods:** Here we use an ethanol exposure paradigm with *nkx2.3* knockdown approaches to test if loss of *nkx2.3* sensitizes Bmp mutants to ethanol induced facial defects. We then combine a morphometric approach with Hybridization Chain Reaction and immunofluorescence to examine the cellular mechanisms underlying Bmp-ethanol interactions.

**Results:** We show that Bmp-ethanol interactions alter morphology of the endodermal pouches, independent of *nkx2.3* gene expression. Morpholino knock down of *nkx2.3* does not sensitize wild type or *bmp4* mutant larvae to ethanol-induced facial defects. However, we did observe a significant increase CNCC apoptosis in ethanol-treated Bmp mutants.

**Conclusions:** Collectively, our results suggest that ethanol’s mode of action is independent of downstream Bmp effectors, converging on CNCC cell survival. Ultimately, our work provides a mechanistic paradigm of ethanol-induced facial defects and connects ethanol exposure with concrete cellular events.

## Introduction

Fetal Alcohol Spectrum Disorders (FASD) is a major issue in the US, impacting up to 5% of live births, making prenatal alcohol exposure (PAE) the leading preventable cause of birth defects resulting in severe societal impact [1, 2]. FASD is an umbrella term describing a range of possible diagnoses for PAE individuals, amongst which FAS is at the most severe end of the spectrum, encompassing a wide range of adverse effects including defects to the facial skeleton [1]. Hallmark facial features of PAE include reduced palpebral fissure length, smooth philtrum, thin upper vermillion lip border and, for this manuscript, underdeveloped jaw [1]. Ethanol dosage and timing and genetic predisposition contribute to FASD etiology [1, 2]. However, genetic predisposition and, more importantly, the cellular and molecular events they regulate, that give rise to the ethanol-induced facial defects in FASD, are still poorly understood.

Zebrafish is an excellent animal model to study gene-ethanol interactions due to its genetic tractability, external fertilization, translucent embryos. In addition, external fertilization allows us to precisely control the timing and dosage of ethanol [3]. Zebrafish embryos develop rapidly with craniofacial skeleton able to be labeled and visualized by 4-5 days post fertilization (dpf). Genetic manipulation with powerful techniques such as CRISPR/Cas mutagenesis and Tol2 transgenesis allows us to readily generate mutant and transgenic lines, respectively. Due to its high fecundity, zebrafish studies can easily generate hundreds of these genetically manipulated embryos enabling high-throughput analysis. This, combined with easy visualization of the embryos using 4D confocal imaging techniques, demonstrate that zebrafish can quite readily generate a large volume of meaningful data. Moreover, due to genetic conservation, these studies will be directly applicable to human since over 70% of genes in zebrafish have homologs in human, with this percentage increasing to 82% when the genes are known to cause human diseases [3, 4]. Therefore, zebrafish is an ideal model in which to study genetic and environmental factors, as well as their interactions, in craniofacial malformations.

In all vertebrates, the craniofacial skeleton is derived from cranial neural crest cells (CNCC), which originate from the dorsal neural tube and migrate and condense in the pharyngeal arches [3, 5, 6]. However, proper facial development can only be achieved by complex interactions between the CNCCs and the surrounding facial epithelia, in particular the pharyngeal endoderm [5]. While the CNCCs are undergoing their migration, cells of the pharyngeal endoderm also undergo their own morphogenesis to form lateral protrusions called pouches [6]. This pouch-forming process involves subsets of endodermal cells reorganizing their points of cell adhesion and following a migratory cue to out-pocket and form the pouches [7, 8]. These pouches then segment the pharyngeal arches and provide scaffolding as well as signaling centers that guide the CNCCs throughout facial development [9]. Perturbations to pouch morphogenesis disrupts their signaling interactions with the CNCCs, resulting in defects to the facial skeleton [6–8]. Together, the endodermal pouches and their interactions with the CNCCs are indispensable for craniofacial development [6, 10].

Multiple genetic signaling pathways have been shown to regulate pouch morphogenesis, including the Bone Morphogenetic Protein (Bmp) signaling pathway [2, 6, 8, 11, 12]. Bmp signaling regulates a wide range of developmental processes, including endodermal pouch morphogenesis [5, 6]. We have previously shown that Bmp signaling is critical for the pouch formation, with disruption in pouch development leading to severe craniofacial defects [6]. Inhibition of Bmp signaling with the small chemical inhibitor, Dorsomorphin (DM) from 10-18 hpf reduces Fgf signaling responses in the endoderm by regulating the expression *fgfr4* [6]. Previous work has also shown that Bmp signaling is also required for specification of pre-pouch endoderm through the expression of *nkx2.3*, and that loss *nkx2.3* disrupts formation of the posterior craniofacial cartilages, in part by regulating expression of the Fgf ligands, *fgf3* and *fgf8a* [11, 13]. Examination of the timing of gene expression in both wild type and Bmp knock down showed that *nkx2.3* is expressed and impacted by Bmp signaling loss earlier that Fgf signaling placing *nkx2.3* upstream of Fgf in pouch development [6, 11, 13]. Overall, this suggests that a complex Bmp-*nkx2.3* signaling pathway regulates pouch development.

We have recently shown that multiple Bmp mutants are sensitive to ethanol-induced facial defects when exposed during DM sensitive time window, 10-18 hpf [14]. These Bmp-ethanol interactions increase the size of the anterior endoderm, disrupting *fgf8a* expression in the oral ectoderm, leading to jaw defects [15]. In addition to jaw defects, we observed defects to the posterior cartilage elements in ethanol-treated Bmp mutants [15]. Strikingly, we observed that Bmp signaling responses are not sensitive to ethanol [15]. This suggests that the interaction between Bmp signaling and ethanol disrupts endoderm morphogenesis downstream of Bmp signaling, potentially interacting with downstream targets to Bmp signaling, such as *nkx2.3*. However, the expression of *nkx2.3* in ethanol-treated Bmp mutants and its role in ethanol-induced pouch and facial defects in Bmp mutants remain unanswered.

Here, we investigate the Bmp downstream target, *nkx2.3*, to determine the mechanism of ethanol action on pouch morphogenesis and subsequent craniofacial development. We show that ethanol does not alter the expression of *nkx2.3* in wild type or *bmp4; smad5* double mutant embryos. We go on to show that *nkx2.3* morphants are not ethanol sensitive and do not further sensitize *bmp4* single mutant embryos to ethanol-induced craniofacial defects. This suggests that ethanol is acting entirely independent of Bmp signaling and its downstream target, *nkx2.3*. Previous work has shown that the CNCCs are sensitive to ethanol-induced apoptosis shifting our analyses, where we show that our Bmp-ethanol interaction significantly increases apoptosis in the CNCCs over all loss of Bmp signaling or ethanol treatment alone. Ultimately, this suggests a two-hit model where loss of Bmp signaling alters endoderm morphogenesis which sensitizes embryos to ethanol-induced CNCCs apoptosis leading to craniofacial defects.

## Materials and Methods

### Zebrafish (Danio rerio) care and use

Zebrafish embryos were raised and cared for using IACUC protocols approved by the University of Louisville. Adult fish were maintained at 28.5 °C. with a 14/ 10-hour light/dark cycle. The *bmp4^st72^* [16], *smad5^b1100^* [17], *sox17:EGFP^s870^* [18], and *sox10:EGFP^ba2^* [19] zebrafish lines were previously described.

### Zebrafish staging and ethanol treatment

Eggs from random heterozygous crosses were collected and embryos were morphologically staged [20], sorted into sample groups of 100 and reared at 28.5 °C to desired developmental time points. All groups were incubated in embryo media (EM). At 10 hpf, EM was changed to either fresh EM or EM containing 1% ethanol (v/v). At 18 hpf, EM containing ethanol was washed out with 3 fresh changes of EM

### Hybridization Chain Reaction (HCR), and immunofluorescence

Embryos were collected at 18 hpf, dechorionated and fixed in 4% paraformaldehyde/PBS overnight at 4°C. HCR protocol was previously described [21]. HCR amplifiers and buffers were acquired from Molecular Instruments. HCR probes against *nkx2.3* were designed as previously described [22]. Immunofluorescence was performed as previously described [6]. Primary antibody against Cleaved Caspase-3 (#9661, Cell Signaling) was used at 1/200. Secondary antibodies Alexa Fluor anti-rabbit 568 was used at 1/500. Confocal images were taken using an Olympus FV1000 or an Olympus FV3000 microscope.

### Pouch measurements

Linear measures of the pouches were obtained in FIJI [23] from 36 hpf embryos. The length of the pouch was defined as the distance from the most dorsal initial branching point from the medial endoderm to the most ventral end of the out-pocketing endoderm. The width of the pouch was reported as the average of three measurements in the medial portion of each pouch, equidistance from the most dorsal and most ventral end of each pouch. Pouch depth was determined using orthogonal view (Z- axis) of each pouch and recorded from where the medial endoderm is no longer visible to the most lateral tip. We measured pouches 1-4 individually, but pouches 5 and 6 were not clearly separated and were measured as a combined single pouch. Volumetric measurements were measured using FIJI.

From each endoderm-labelled confocal stack, we generated a sub-stack for each pouch defining the stack range from where it no longer connects to the medial endoderm to the most lateral tip of the pouch. The image threshold function is then used to determine and record the lower and upper fluorescent threshold of the pixels in the sub-stack. 3D ROI Manager is then used to track the continuous pixels present in each sub-stack. From this, the volume value is measured and recorded as pixel volume (voxel), then converted into microns.

### Morpholino injection and cartilage staining

Wild type or *bmp4* mutant embryos were injected with ∼18 ng of translation blocking *nkx2.*3-MO (Gene Tools) at the 1- to 2-cell stage and raised to 5 dpf. At 5 dpf, zebrafish larva were fixed and facial cartilages were stained with alcian blue [24]. Whole mount, ventral view, brightfield images of the viscerocranium were taken on an Olympus BX53 compound microscope.

### Statistical analyses

Angle of the Ceratohyal (AOC), endoderm measures and cell counts of apoptotic CNCCs were analyzed with a two-way ANOVA (type III) with a Tukey’s Multiple comparisons Test in Graphpad Prism 9.5.1 (Graphpad Software Inc., La Jolla, CA).

## Results

### The pharyngeal pouches are malformed in ethanol-treated Bmp mutants

We have previously shown that mutation in Bmp signaling components sensitizes zebrafish embryos to ethanol-induced jaw malformations through increases in size of the anterior pharyngeal endoderm [15]. Defects to anterior endoderm development are known to impact jaw development but not development of the posterior endoderm cartilage elements, while defects in pouch morphogenesis disrupt formation of the posterior cartilage elements [6, 10]. Our work demonstrated that ethanol broadly disrupts development of the craniofacial skeleton, both the jaw and posterior cartilage elements in ethanol-treated Bmp mutants [15], suggesting that Bmp-ethanol interactions are disrupting pouch morphogenesis in addition to defects to anterior endoderm morphogenesis. Similar to the anterior endoderm, we observed that while the pouches largely form in both Bmp mutants and ethanol-treated embryos their shape and size may be altered (Fig. 1A-D).

**Figure 1:**
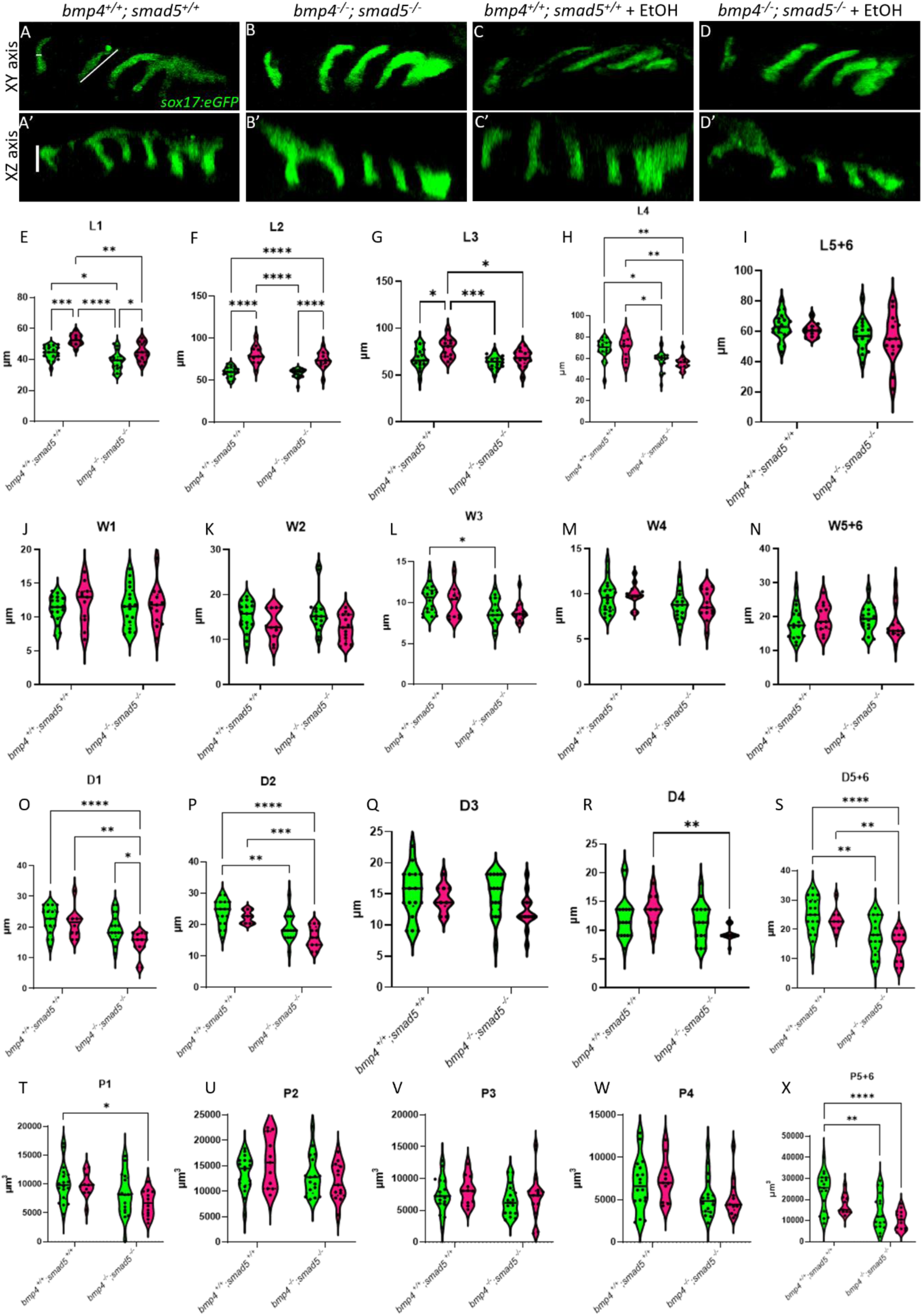
Ethanol alters shape and size of the pharyngeal pouches. (A-D) Representative confocal images at 36 hpf showing the pharyngeal endoderm labeled with *sox17:eGPF* (lateral view, anterior to the left). Pouches 1-4 are shown, pouches 5 and 6 have yet to fully separate and are counted as a single pouch (5+6). The length was measured from the start to the end of the out-pocketing endoderm. The width of the pouch was measured in the medial portion of that specified pouch, perpendicular to the length. Z depth was determined using orthogonal view (XZ) and recorded from the stack where the medial endoderm of is not visible to the tip of the pouch. (E-S) Linear measurements of each pharyngeal pouch. (T-X) Measures of pouch volume. *: P≤0.05. **: P≤0.01. Every additional “*” reduces the P value by a factor of 10.

To test this, we analyzed the length, width, z-depth, and volume of the pharyngeal pouches in ethanol- treated *bmp4; smad5* double mutant embryos (defined as “Bmp mutants” henceforth) at 36 hpf. We have previously shown that this double mutant line is 100% penetrant for highly expressive ethanol- induced defects to the facial skeleton making [15]. Pouch length was defined as the distance from the dorsal-most end to the ventral-most end of the pouch (Fig. 1A); pouch width was measured equidistance from the pouch ends, perpendicular to the length (Fig. 1A); and the z depth was determined using orthogonal view and recorded from the stack where the medial endoderm of is not visible to the tip of the pouch (Fig. 1A’). We measured pouches 1-4 individually, but pouches 5 and 6 were not clearly separated and were measured as a combined single pouch.

Overall, pouch formation was not largely disrupted due to genotype or treatment. However, while variable between the pouches, we did find significant changes in pouch length and depth while pouch width was largely unaffected (Fig. 1E-W). Pouches 1, 2 and 3 were significantly elongated by genotype, ethanol-treatment and genotype by ethanol treatment (Fig. 1E-G). Pouch 4 showed only significant elongation due to genotype, and pouch 5+6 are largely unaffected (Fig. 1H-I). For pouch depth, pouch 1 shows a significant reduction in depth in ethanol-treated Bmp mutants compared to untreated wild type and Bmp mutant and ethanol-treated wild type siblings (Fig. 1O). Pouches 2, 5+6 show a reduction in depth in Bmp mutants compared to wild type siblings, independent from treatment (Fig. 1P,S). Pouches 3 and 4 were largely unaffected (Fig. 1Q,R). Finally, measuring the volume of each individual pouch, only the pouches 1 and 5+6 show significant reduction in volume in ethanol-treated mutants (Fig. 1T,X). Overall, these measurements indicate that while the pouches largely form, Bmp-ethanol interactions alter size and shape of the pouches suggesting that these changes in pouch shape and size may be impacting facial development in ethanol-treated Bmp mutants

### Expression of *nkx2.3* is reduced in ethanol-treated Bmp mutants

We have previously shown that ethanol does not attenuate Bmp signaling [15] suggesting that ethanol may be interacting with downstream Bmp targets to disrupt craniofacial development. Work from Li et al., (2019) showed that *nkx2.3* is a downstream target of Bmp signaling regulating pouch specification [11]. To test whether ethanol affects downstream Bmp targets, we used hybridization chain reaction (HCR) to assess the expression of *nkx2.3* at 18 hpf in wild type and Bmp mutant embryos with the endoderm labeled with the *sox17:eGFP* transgene. At 18 hpf, *nkx2.3* is expressed most in the developing pouch and ethanol does not reduce this expression (Fig. 2A,C). Bmp mutant embryos show reduced *nkx2.3* expression regardless of ethanol (Fig. 2B,D). Volumetric measurements of colocalized fluorescent intensity of *nkx2.3* and *sox17:eGFP* expression in the pouches in the untreated and ethanol-treated wild type and untreated and ethanol-treated Bmp mutants are 13781, 10974, 8646, and 5726 µm^3^, respectively (Fig. 2E). Despite the statistically significant differences between untreated wild-type and ethanol-treated Bmp mutant embryos observed in *nkx2.3* colocalization in pouches, our data demonstrates that ethanol does not directly attenuate *nkx2.3* expression in the endoderm in wild type or Bmp mutant embryos (Fig. 2E). One possible explanation for this significant difference between untreated wild type and ethanol-treated mutant embryos is the difference in variation observed in untreated wild type vs ethanol-treated Bmp mutant embryos (Fig. 2E), though further investigation is needed. Despite this, these data overall suggest that ethanol does not attenuate *nkx2.3* expression at the transcription level.

**Figure 2:**
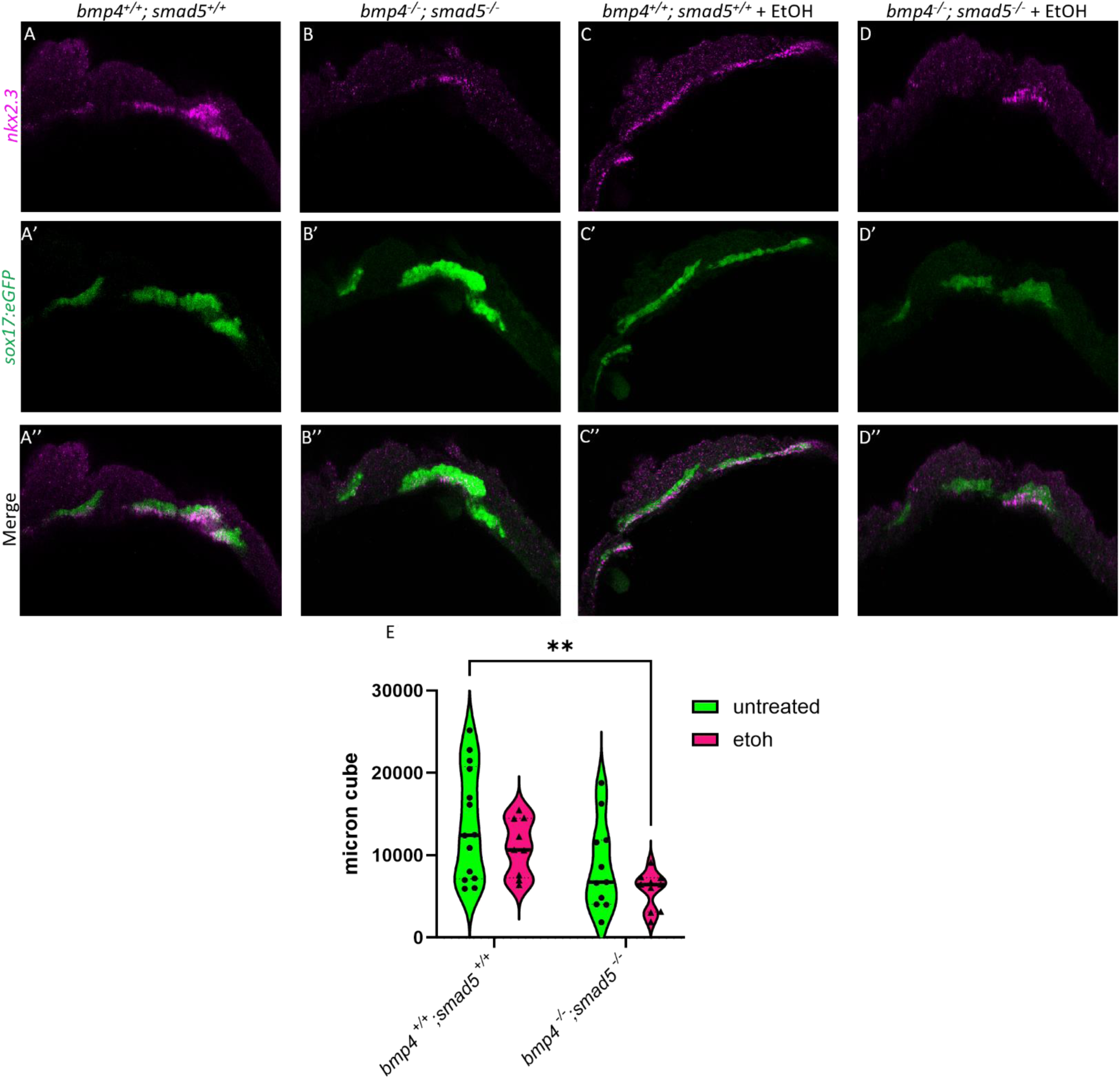
Ethanol does not alter *nkx2.3* expression in Bmp mutants. (A-D”) Confocal images at of sox17:eGPF labeled endoderm (green) and *nkx2.3* mRNA (magenta) expression using HCR at 18 hpf (lateral views, anterior to the left). (E) Volume measurements of colocalized intensity of *nkx2.3* gene expression and the out-pocketing pouch. Solid lines indicates the median of that treatment group. Dash lines show 95% confidence of the median. **: P≤0.01.

### Ethanol does not sensitize *nkx2.3* morphants to craniofacial defects

While ethanol does not decrease *nkx2.3* gene expression, it may still interact with the *nkx2.3* gene product. Loss of *nkx2.3* does lead to defects to the posterior cartilage elements [13]. To test if *nkx2.3* knock down sensitizes embryos to ethanol-induced facial defects, we injected wild type embryos with *nkx2.3* translation blocking morpholino and treated them with ethanol from 10-18 hpf, when *nkx2.3* is required for endoderm specification [11]. Knockdown of *nkx2.3* resulted in a range of phenotypes from loss of the ceratobranchials to inversion of the ceratohyal. We categorize these phenotypes based on the severity of facial defects (Fig. 3A-D). Category 1 are wild type in phenotype with the angle of ceratohyal (AOC) less than 90° (Fig. 3A). Category 2 includes mild phenotype of AOC increased compared to wild type larvae, ranging from 90 and 135° (Fig. 3B). Category 3 are those larvae with the AOC ranging from 135 to 180° (Fig. 3C). Finally, category 4 are larvae where the AOC is more than 180°. Category 4 larvae can also display loss of jaw, ceratohyal and / or ceratobranchial (CB) loss (Fig. 3D). AOC is significantly increased in *nkx2.3* knock down but is not impacted by ethanol treatment (Fig. 3E). Non-injected larvae are normal phenotypically regardless of ethanol treatment (100% vs 93.5%). Larvae injected with *nkx2.3* MO have a lower percentage of larvae in category 1, 45.95%, and increased percentage in other categories: category 2: 13.5%, category 3: 10.2%, and category 4: 29.73% (Table 1). Consistent with our AOC measures, ethanol does not increase penetrance of facial defects in *nkx2.3* morphants (Table 1), indicating an insensitivity of *nkx2.3-*knocked down embryos to ethanol.

**Figure 3:**
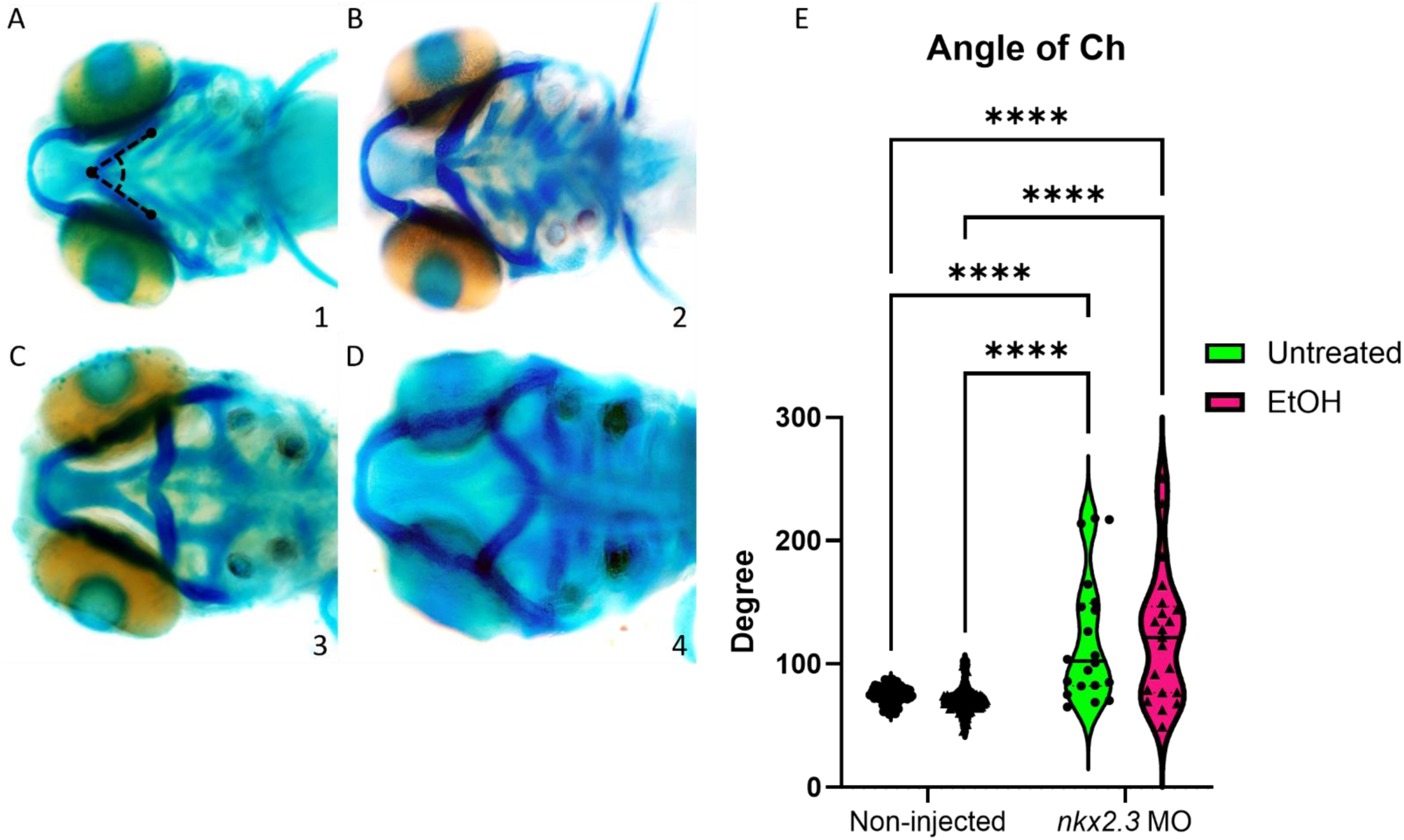
Ethanol does not exacerbate facial defects in *nkx2.3* morphants. (A-D) 5 dpf alcian- stained larvae non-injected and *nkx2.3*-MO injected (∼18 ng; ventral views, anterior to the left). 4 categories were used to assess the severity of facial defects. Category #1 consists of larvae with the angle of ceratohyal (AOE) < 90°. Category #2 are those that have the AOE from 90-135°. Category #3 are those ranging from 135 and 180. Category #4 are embryos with severe defects, including those with either inverted Ch (AOC > 180), jaw loss, or Ch/Cb loss. (E) Graph of AOC of untreated and ethanol-treated non-injected larvae and *nkx2.3* morphants. Solid lines indicate the median of that treatment group. Dash lines show 95% confidence of the median. ****: P≤0.0001

**Table 1:** Penetrance and expressivity of *nkx2.3* MO injected wild type larvae in ethanol. Percent of larvae, either non-injected or *nkx2.3* MO injected, that show the phenotypes in Fig. 3 for each treatment group. The numbers in parentheses in different phenotype categories show the number of larvae overall and those exhibiting each phenotype category.

| Table 1: Ethanol-*nkx2.3* penetrance and expressivity |  |  |  |  |  |
| --- | --- | --- | --- | --- | --- |
|  |  | Phenotype categories |  |  |  |
|  |  | 1 | 2 | 3 | 4 |
| Control | Non-injected (87) | 100% (87) | 0 | 0 | 0 |
|  | *nkx2.3* MO (37) | 45.95% (17) | 13.5% (5) | 10.2% (4) | 29.73% (11) |
| 1% EtOH | Non-injected (77) | 93.51% (72) | 6.49% (5) | 0 | 0 |
|  | *nkx2.3* MO (57) | 57.89% (33) | 8.77% (5) | 3.51% (2) | 29.82% (17) |

### Ethanol does not exacerbate facial defects *bmp4* mutants with *nkx2.3* MO

Larvae lacking *bmp4* are ethanol sensitive and exhibit facial defects [15]. We hypothesized that knocking down *nkx2.3* in *bmp4* mutant larvae would further sensitizes these larvae to ethanol, exacerbating their ethanol-induced phenotypes closer to the *bmp4;smad5* double mutant and dorsomorphin-treated larvae [6, 15]. Similarly to the *nkx2.3* MO injected wild type larvae, we categorized the facial phenotype into 4 groups based on the AOC and presence of facial components as mentioned above. As observed in Fig. 3, ethanol treatment in wild type larvae injected with the *nkx2.3* MO does not significantly change the AOC or the distribution of the percentage of phenotypes nor (Fig. 4A-E, Table 2). Morpholino knockdown of *nkx2.3* in ethanol-treated *bmp4* mutant larvae shifts the phenotype categories compared to untreated *nkx2.3-*MO; *bmp4* mutant larvae (56.67% to 28.57% in category 1 and 6.67% to 28.57% in category 2, Table 2). However, this mild disruption of the facial skeleton was observed previously in ethanol-treated *bmp4* mutants, suggesting that this result is simply due to ethanol sensitivity in *bmp4* mutant larvae [15]. Surprisingly, no significant increase in AOC or phenotype category 3 and 4 was detected in ethanol-treated *nkx2.3*-MO; *bmp4* mutant larvae (Fig. 4E, Table 2). These data indicate that knocking down *nkx2.3* in *bmp4* mutant larvae does not exacerbate the ethanol-induced facial defects, suggesting that ethanol is acting independent of Bmp signaling to disrupt facial development.

**Figure 4:**
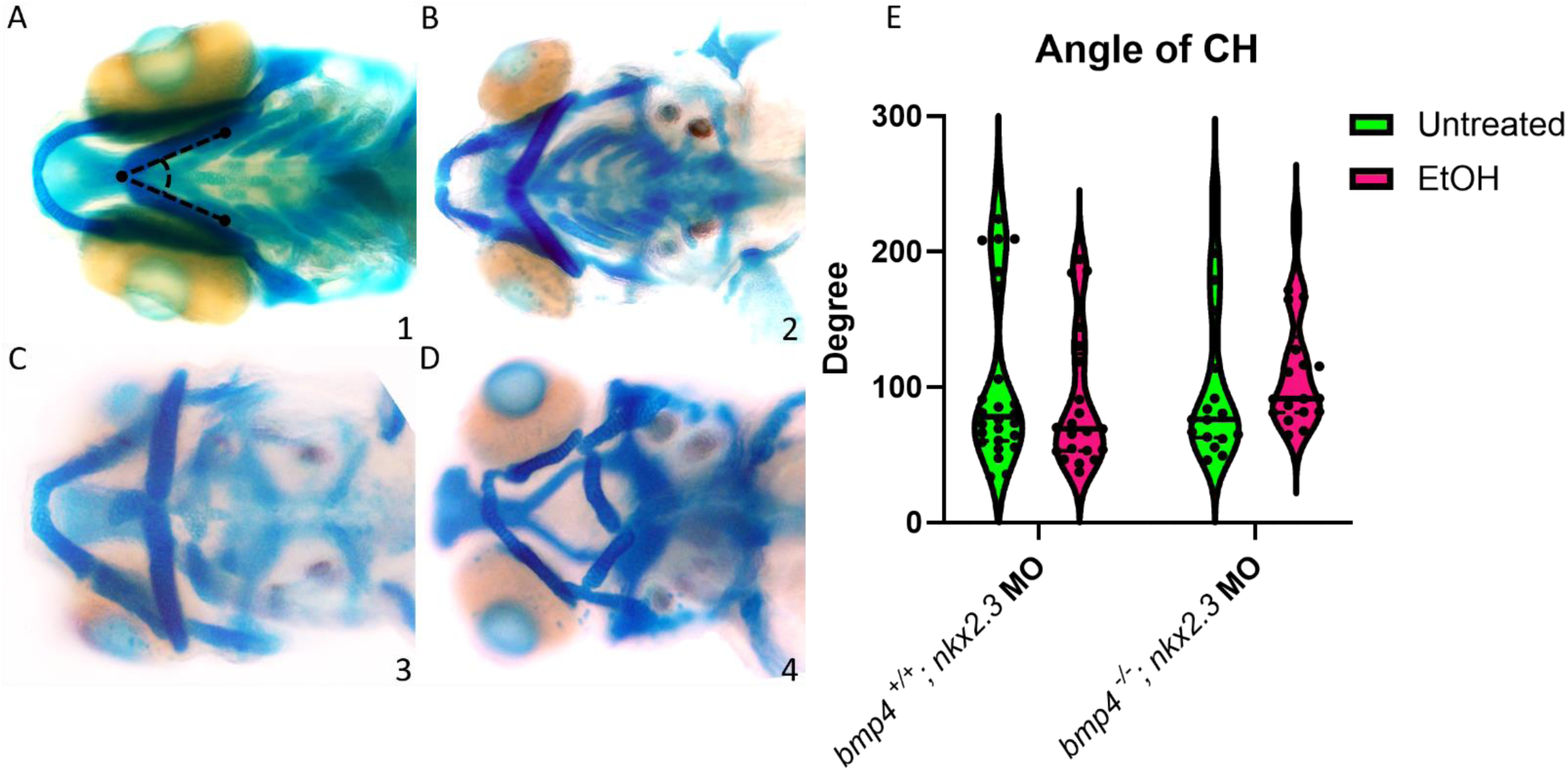
K**n**ockdown **of *nkx2.3* does not further sensitize the in *bmp4* mutant larvae to ethanol.** (A-D) 5 dpf alcian-stained larvae untreated and ethanol-treated, *nkx2.3*-MO injected (∼18 ng) wild type and *bmp4* mutant larvae (ventral view, anterior to the left). 4 categories were used to assess the severity of facial defects. Category #1 consists of larvae with the angle of ceratohyal (AOE) < 90°. Category #2 are those that have the AOE from 90-135°. Category #3 are those ranging from 135 and 180. Category #4 are embryos with severe defects, including those with either inverted Ch (AOC > 180), jaw loss, or Ch/Cb loss. (E) Graph of AOC of untreated and ethanol-treated, *nkx2.3*-MO injected wild type and *bmp4* mutant larvae. Solid lines indicate the median of that treatment group. Dash lines show 95% confidence of the median. ****: P≤0.0001

**Table 2:** Penetrance and expressivity of *nkx2.3* MO injected wild type and *bmp4* mutant larvae in ethanol. Percent of *nkx2.3* MO injected wild type or *bmp4* mutant larvae, that show the phenotypes in Fig. 4 for each treatment group. The numbers in parentheses in different phenotype categories show the number of larvae overall and those exhibiting each phenotype category.

| Table 2: Ethanol-*nkx2.3*-*bmp4* penetrance and expressivity |  |  |  |  |  |
| --- | --- | --- | --- | --- | --- |
|  |  | Phenotype categories |  |  |  |
|  |  | 1 | 2 | 3 | 4 |
| Control | *bmp4*<sup>+/+</sup> (30) | 56.67% (17) | 6.67% (2) | 3.33% (1) | 33.33% (10) |
|  | *bmp4*<sup>-/-</sup> (19) | 57.89% (11) | 10.53% (2) | 0 (0) | 31.58% (6) |
| 1% EtOH | *bmp4*<sup>+/+</sup> (21) | 66.67% (14) | 4.76% (1) | 0 (0) | 28.57% (6) |
|  | *bmp4*<sup>-/-</sup> (21) | 28.57% (6) | 28.57% (6) | 4.76% (1) | 38.09% (8) |

### Ethanol sensitizes Bmp mutants to increased CNCCs apoptosis

The endoderm is critical in multiple aspects of craniofacial development, in particular CNCC survival [25, 26]. In embryos that completely lack endoderm, *sox32* (*cas*) mutants, CNCCs migrate and condense in the pharyngeal arches but undergo apoptosis contributing to almost complete loss of the facial skeleton [25]. In mutants in endoderm development where the endoderm is present but fails to form properly, CNCC cell death is still observed [8, 27]. In addition, multiple studies have shown that ethanol also induces CNCCs apoptosis in different animal models [28–31], suggesting that ethanol-treated Bmp mutants will have increased CNCC apoptosis. To test that ethanol exacerbates CNCCs apoptosis in Bmp mutants, we performed immunostaining against cleaved Caspase-3 on untreated and ethanol-treated wild type and Bmp mutant embryos labeling the CNCCs with *sox10:eGFP*. The number of apoptotic CNCCs in untreated wild type embryos is low and contained mostly in the anterior first arch (Fig. 5A-A’’). In the untreated Bmp mutant and ethanol-treated wild type embryos, apoptotic CNCCs expand to the posterior of arch 1 and into arch 2 (Fig. 5B-C’’). The number of apoptotic CNCCs in the untreated wild type and Bmp mutant embryos are relatively low with the mean of 7.18 and 9.27 cells, respectively, while the number of apoptotic cells in ethanol-treated wild type embryos increases to a mean of 25.18 cells (Fig. 5E). However, the average number of apoptotic CNCCs in ethanol-treated Bmp mutant is 114.8 cells, significantly increased compared to all other groups, and distributed evenly throughout most of arches 1 and 2 (Fig. 5D). These data indicate that mutation in the Bmp pathway sensitizes CNCCs to ethanol-induced apoptosis and suggests a two-hit model where Bmp mutation in combination with ethanol exposure disrupt endoderm morphogenesis and increases CNCCs apoptosis, leading to facial defects to the facial skeleton.

**Figure 5:**
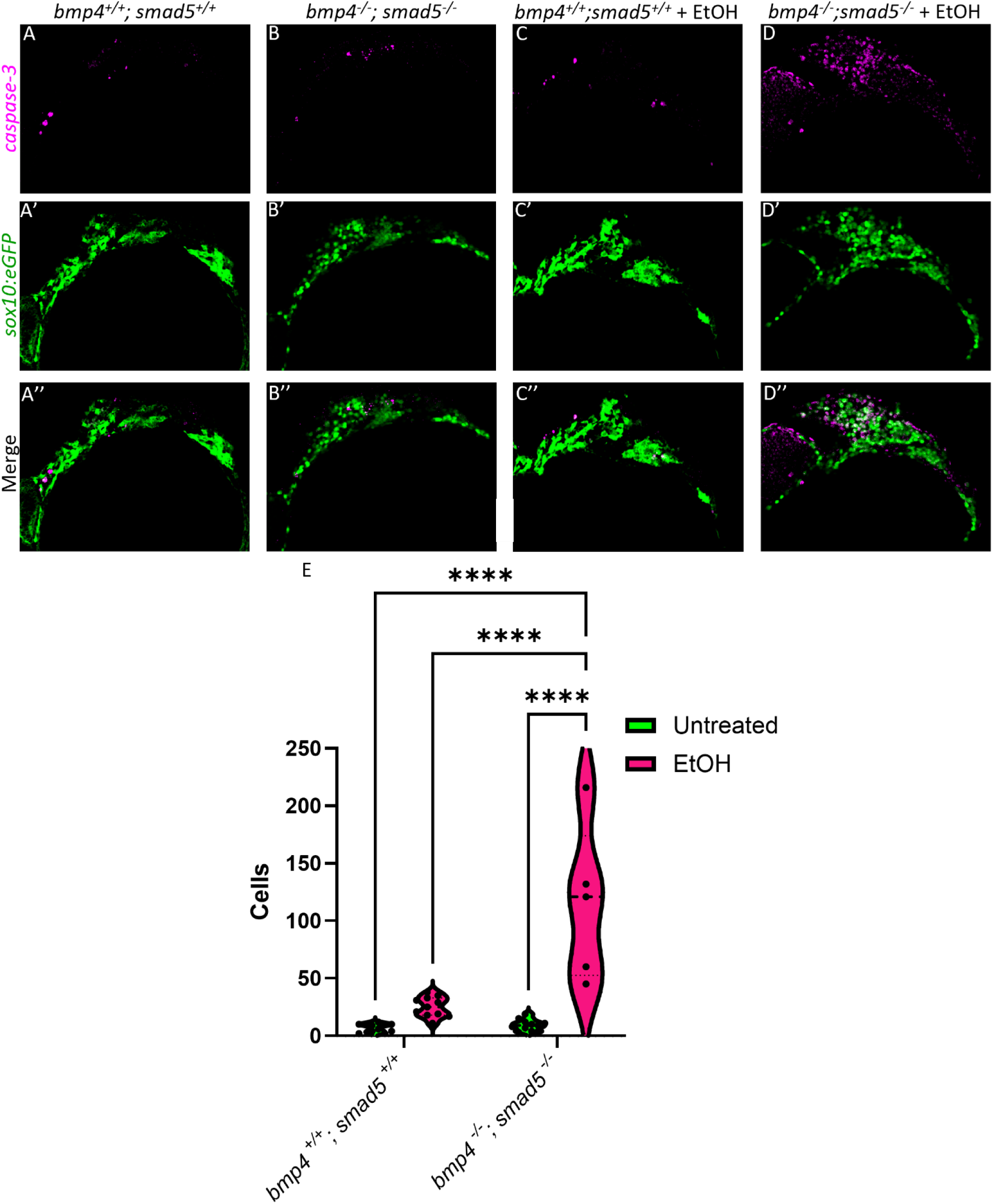
Ethanol-treated Bmp mutants have elevated CNCCs apoptosis. (A-D”) Confocal images of CNCCs (*sox10:eGFP*) and Cleaved Caspase-3 at 18 hpf (lateral view, anterior to the left). (E) Number of detectable apoptotic CNCCs in all arches in untreated and ethanol-treated wild type and Bmp mutant embryos. Solid lines indicate the median of that treatment group. Dash lines show 95% confidence of the median. ****: P≤0.0001.

## Discussion

Pouch formation is critical to overall development of the craniofacial skeleton with disruption of pouch out-pocketing leading malformation and / or loss of viscerocranial cartilage elements in the facial skeleton [6–8]. Multiple pathways are required to regulate several key aspects of pouch formation including the Bmp and Fgf signaling pathways [6–8]. Bmp signaling appears to lie at the head of this regulation driving both pouch out-pocketing via expression of *fgfr4* and pouch specification via *nkx2.3* [6, 11]. Timing of *nkx2.3* gene expression and Fgf signaling responses in wild type and Bmp knock down suggest that *nkx2.3* is upstream of Fgf in pouch development [6, 11]. This suggests that Bmp signaling is a key regulator to pouch morphogenesis with loss of Bmp signaling leading to reduced expression of multiple downstream target genes. We have recently shown that mutation to multiple Bmp signaling components sensitize embryos to ethanol induced defects throughout the viscerocranium [15]. We found that these defects are generated in part by disrupting anterior endoderm morphogenesis [15]. However, while the anterior endoderm is necessary for jaw development, it does not contribute to defects to the posterior cartilage elements [10]. Given that we observed posterior cartilage defects in our ethanol-treated Bmp mutants, this suggests that pouch morphogenesis was also impacted by these Bmp-ethanol interactions. Here, we showed that the ethanol-induced facial skeletal defects in Bmp mutants is independent of the Bmp-*nkx2.3* signaling pathway. While we showed ethanol-induced changes in pouch morphology, we showed *nkx2.3* expression is not impacted by ethanol exposure and that knocking down of *nkx2.3* does not further sensitize wild type nor *bmp4* mutants to ethanol-induced facial defects. Strikingly, we observed increased cell apoptosis in CNCCs in ethanol-treated Bmp mutants suggesting a two-hit model where ethanol sensitizes Bmp mutants CNCC cell death leading to facial defects.

### Ethanol induces cell death in Bmp mutants

Our previous study has shown blocking Bmp signaling does not alter endoderm cell numbers [6]. Yet, our data here shows changes in pouch shape, but not overall volume suggesting that ethanol is not changing the number of endoderm cells in any meaningful way. In our previous work, we saw loss of Bmp signaling responses in the endoderm in Bmp mutants regardless of ethanol-treatment, while we did not observe any changes to Bmp signaling responses in non-endoderm tissues either due to ethanol exposure or in Bmp mutants [15]. This demonstrates that ethanol does not directly impinge on Bmp signaling activity raising the question of the mode of action of ethanol on facial development. Multiple lines of evidence show that ethanol induces cell death in the CNCCs [28–31]. Our data shows that our Bmp-ethanol interactions significantly increase cell apoptosis in migrating CNCC. However, we only observed this increase in ethanol-treated Bmp mutants, not in ethanol-treated wild type or untreated Bmp mutant larvae alone, suggesting a two-hit model where Bmp mutation sensitizes embryos to ethanol-induced cell death in the CNCCs, though the exact mechanism remains an enigma. Several mechanisms have been suggested including increase oxidative stress, mTOR mis-regulation and changes in calcium flux [32, 33]. Future work will be needed to determine the mechanism underlying CNCC cell death in our ethanol-sensitive Bmp mutants and how disruption to endoderm morphology contributes.

Our ethanol exposure paradigm is sub-teratogenic only observing increased CNCC apoptosis in ethanol-treated Bmp mutants implicating a role for the endoderm morphogenesis in cell survival in migrating CNCCs. In *sox32* mutants that completely lack endoderm, CNCCs properly migrate and condense in the pharyngeal arches but show increased CNCC cell death after condensation [25]. In mutants that still generate the endoderm but display defects in pouch size and shape, such as *vgll2a*, increased CNCC cell death in the arches is still observed [27]. Combined this data implicates CNCCs receive cell survival cues from the endoderm when in close proximity, either through direct cell-cell contacts or close, paracrine-based cell signaling. We observed similar pouch shape changes in our ethanol-treated Bmp mutants, yet, unlike these previous examples, we observed CNCC apoptosis in migrating CNCCs, some distance from the forming pouches. This suggests that endoderm provides cell survival cues to the migrating CNCCs prior to completed pouch out-pocketing and CNCC condensation in the pharyngeal arches. However, the cell survival signals from the endoderm and how they are transmitted to the CNCCs remain largely unknown. Future work using transcriptomic approaches on endodermal cell will help shed light on these signals.

### Ethanol sensitive Bmp-dependent pathway regulating pouch development

We have previously shown that loss of Bmp signaling result in severe morphological defects to the endoderm through the downstream target Fgf signaling [6]. Work from Li et al., 2019 [11] added *nkx2.3* as another downstream target of Bmp signaling in pouch formation. More recent work from the same lab showed that the pouches largely form properly, though subtle shape changes can still be observed in *nxk2.3* morphants [13]. While not as severe, Bmp-ethanol interactions also disrupt endoderm morphogenesis, though ethanol does not act on Bmp signaling [15] or its downstream target of *nkx2.3* (This study). However, ethanol may still act on additional downstream targets of Bmp signaling beyond *nkx2.3*. It is possible that ethanol may be acting on Fgf signaling leading to defects in endoderm morphogenesis. Fgf signaling acts as the migratory cue to the endoderm for pouch out-pocketing [8]. We have shown that Bmp signaling regulates reception of Fgf signaling through expression of *fgfr4* in the pre-pouch endoderm [6]. Although Fgf mutants have pouch defects similar to Bmp loss [6–8], our pilot screens show they are not ethanol-sensitive though these results need to be confirmed (data not shown) as we are currently exploring the role of Fgf signaling in our Bmp-ethanol interactions.

Another potential Bmp target in pouch development is Wnt/PCP signaling pathway. Multiple Wnt ligands such as *wnt11r* and *wnt4a* had been shown to take part in the pouch morphogenesis through destabilization and re-stabilization of points of cell adhesion necessary to promote pouch out-pocketing [7, 8]. We have observed reductions of expression of *fzd8a* in the endoderm in DM-treated embryos (data not shown) suggesting that Bmp signaling maybe be regulating reception of Wnt/PCP signaling similar to its regulation of Fgf signaling [6]. Moreover, it has been shown that the Wnt/PCP pathway members *gpc4* and *vang2* interact with ethanol disrupting craniofacial development [34]. This suggest that Wnt/PCP pathway components maybe downstream ethanol-sensitive targets in Bmp mutants. It is also possible that more than one of these downstream Bmp targets may be driving ethanol sensitivity in Bmp mutants and future work will be needed to dissect all potential interactions between Bmp, Fgf, Wnt/PCP and ethanol during endoderm morphogenesis.

In conclusion, we show that ethanol alters endodermal pouch morphology and subsequent facial development in ethanol-treated Bmp mutants, independent of *nxk2.3*, resulting increased CNCC apoptosis. This suggests a two-hit model where loss of Bmp signaling alters endoderm morphology which sensitizes embryos to ethanol-induced CNCCs apoptosis leading to craniofacial defects. Thus, our work provides a deeper mechanistic understanding of gene-ethanol interactions on the complex signaling and tissue interactions driving craniofacial development. Ultimately, this will provide a conceptual framework and a mechanistic paradigm of ethanol-induced birth defects and connect ethanol exposure with concrete cellular events that could be sensitive beyond facial development.

## Acknowledgements

The authors would like to thank Kevin Kump for zebrafish animal care and husbandry. We also thank Dr. Duygu Özpolat and Dr. Ryan Hull for providing script for HCR probe design.

## Author Contributions

CBL conceived the project. HV and CBL designed all zebrafish studies. HV conducted experiments, analyzed data and generated all figures. HV wrote the manuscript. HV and CBL reviewed and edited the manuscript. All authors have read and approved the final manuscript.

## Funding

This work was funded by National Institutes of Health/National Institute on Alcohol Abuse (NIH/NIAAA) R00AA023560 and R01AA031043 to CBL.

## Institutional Review Board Statement

The study was conducted according to the guidelines of the Animal Welfare Act and PHS policy on Humane Care and Use of Laboratory Animals of the United States of America and approved by the Institutional Animal Care and Use Committee (IACUC) of The University of Louisville (IACUC 24352 2/8/2024).

## Conflicts of Interest

The authors declare that they have no competing or conflicts of interests.

## Notes

### Competing Interest Statement

The authors have declared no competing interest.

